## Supplementary figures and images for "Sex–specific Single Transcript Level Atlas of Vasopressin and its Receptor (AVPR1a) in the Mouse Brain"

### Supplemental Figure 1

#### Supplementary Figure 1A

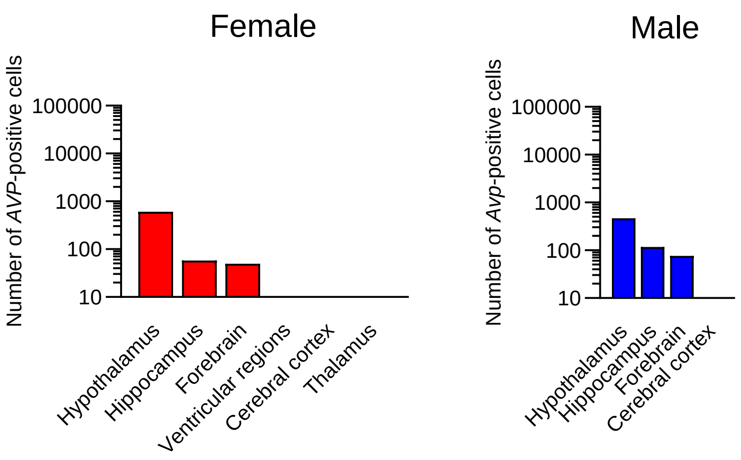

#### Supplementary Figure 1B

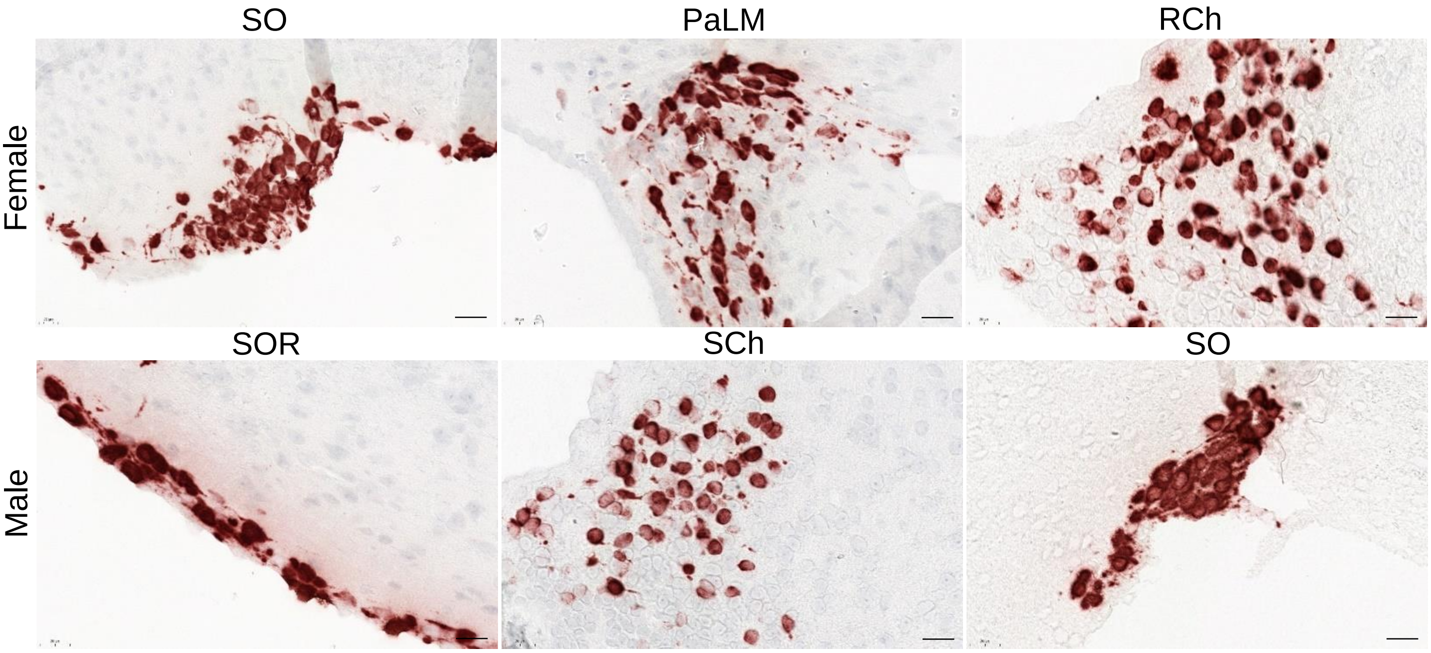

#### Supplementary Figure 1B

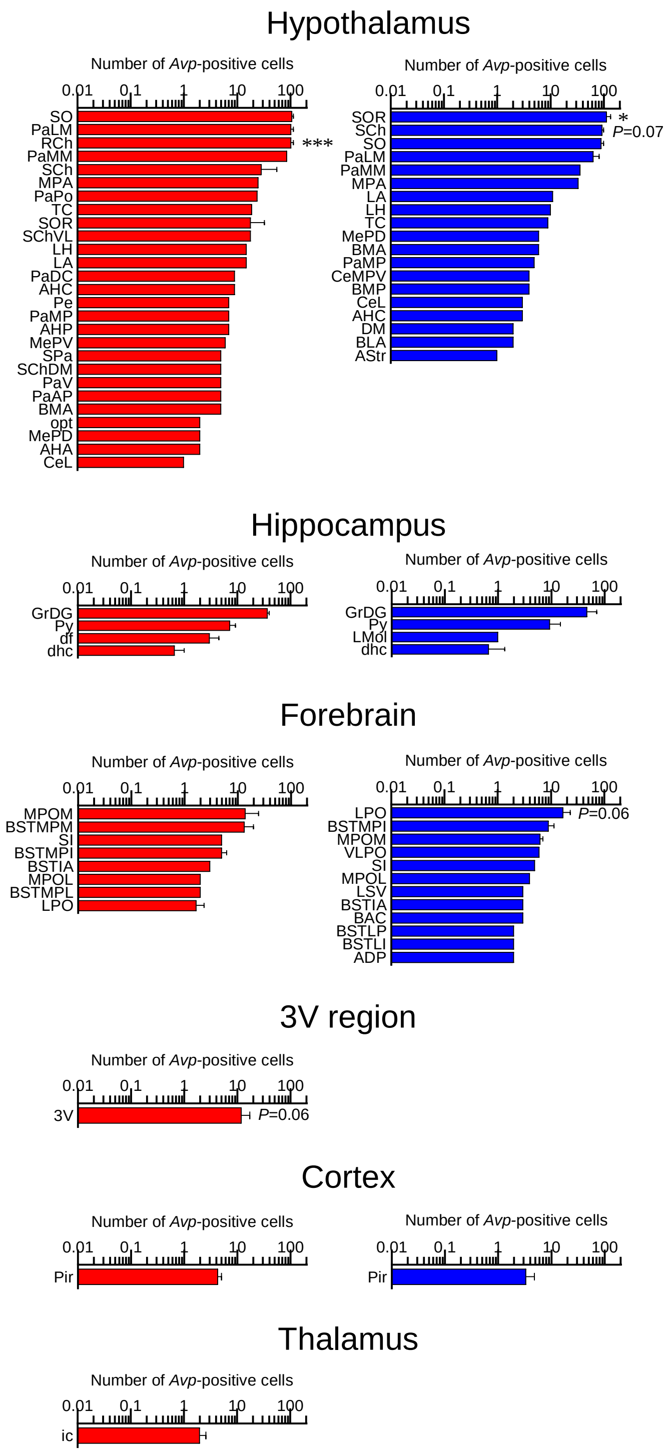
