## Supplemental Figure 2 for "Sex–specific Single Transcript Level Atlas of Vasopressin and its Receptor (AVPR1a) in the Mouse Brain"

### Supplementary Figure 2A

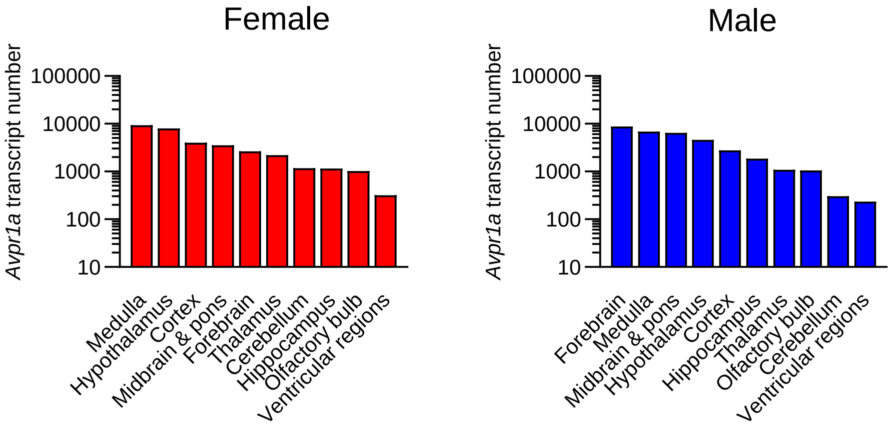

### Supplementary Figure 2B

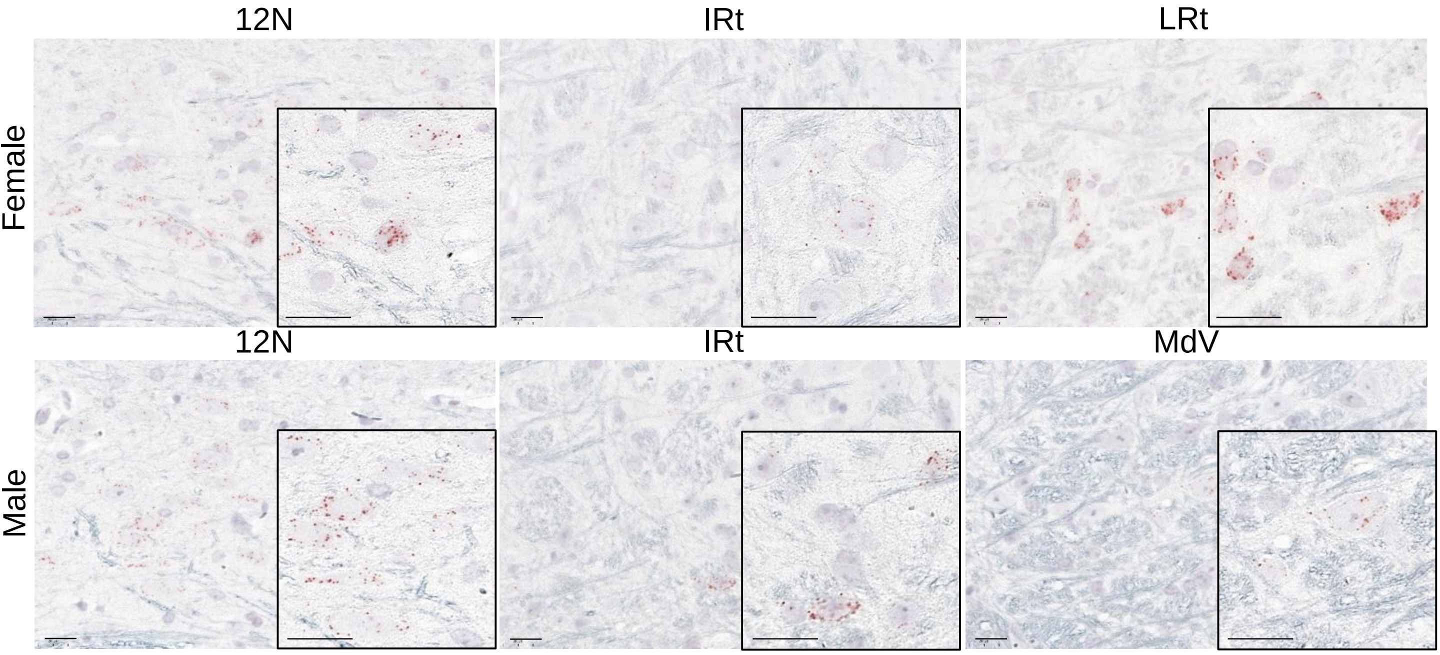

### Supplementary Figure 2B

#### Medulla

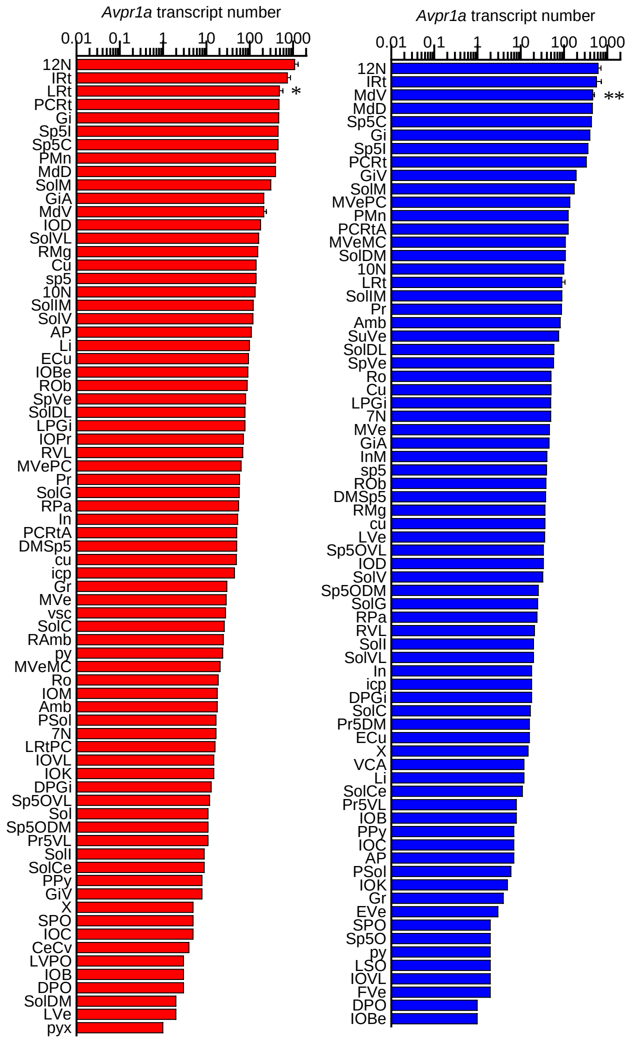

### Supplementary Figure 2B

#### Hypothalamus

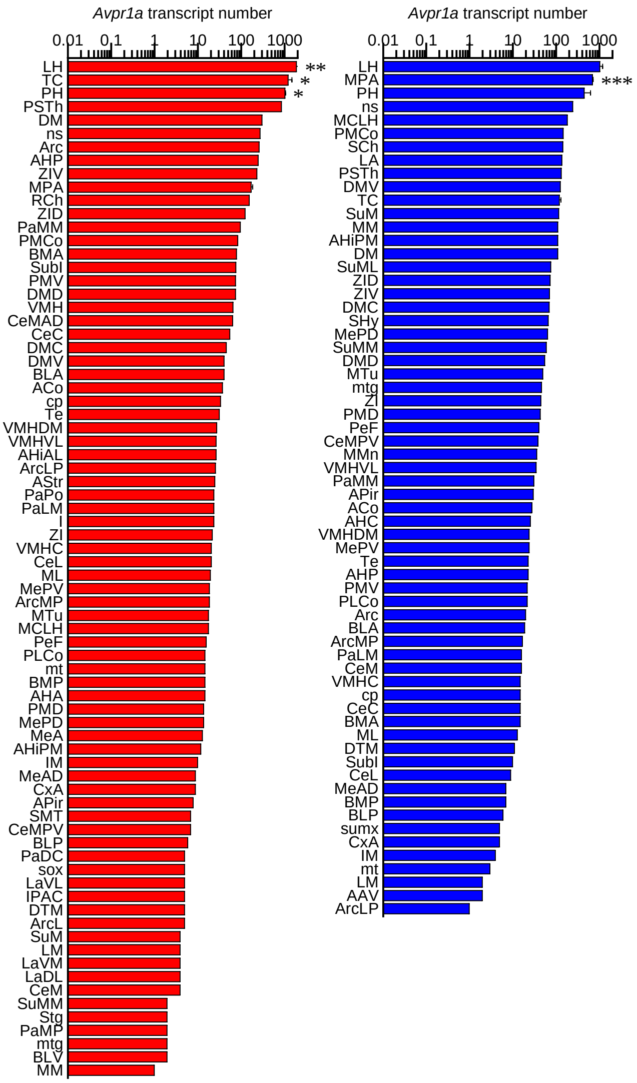


#### Cortex

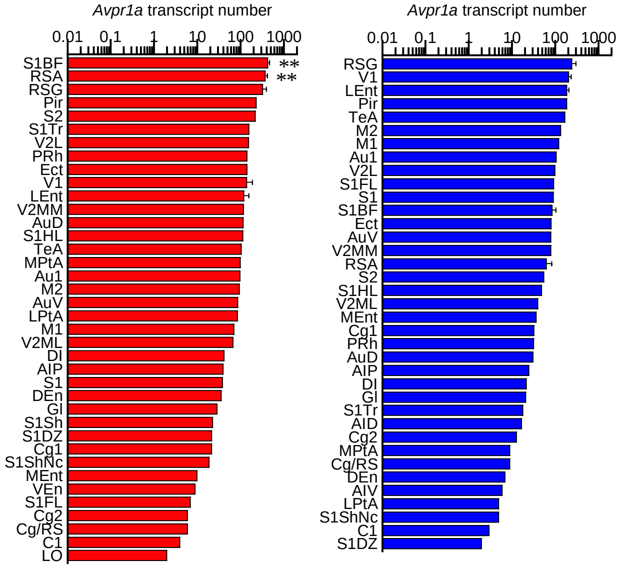

### Supplementary Figure 2B

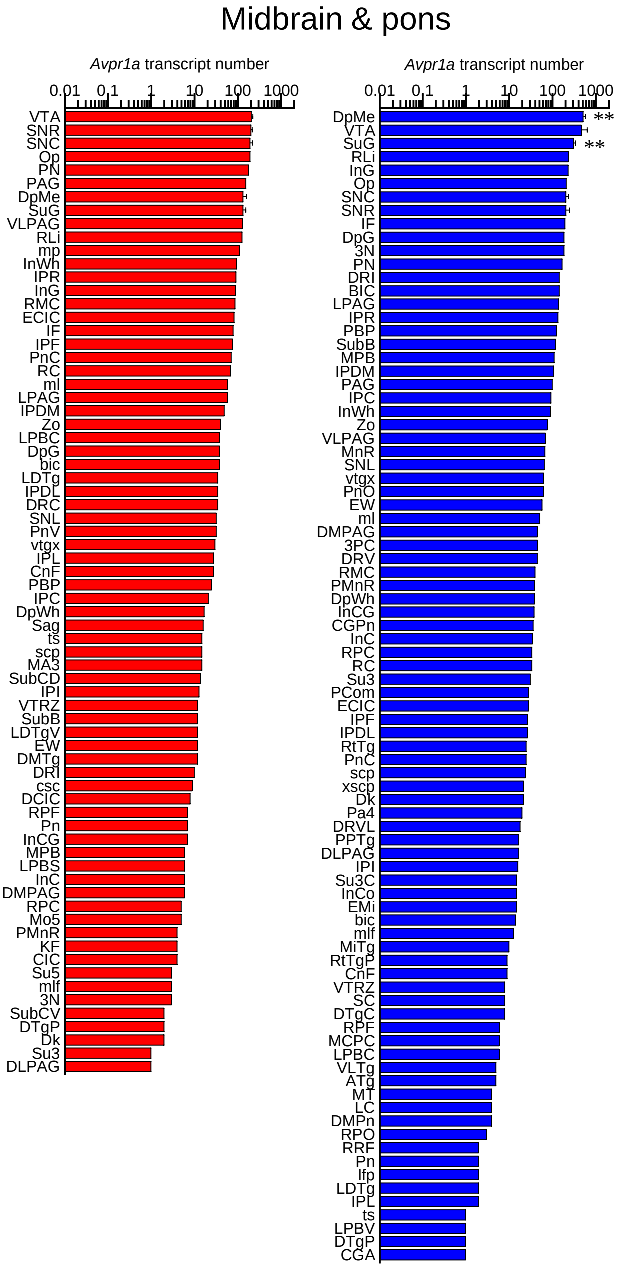

### Supplementary Figure 2B

#### Forebrain

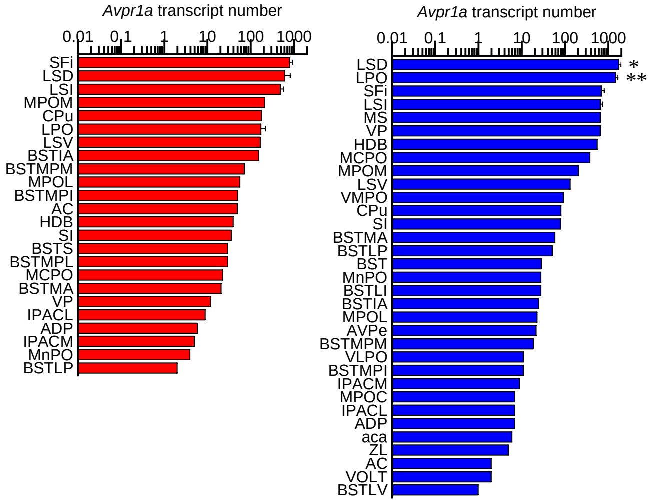


#### Thalamus

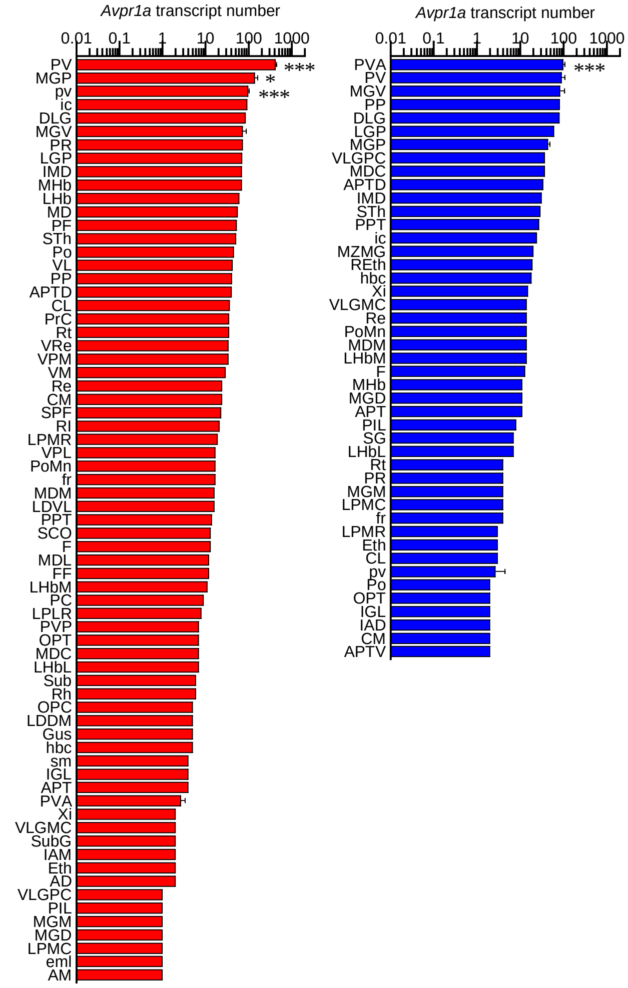

### Supplementary Figure 2B

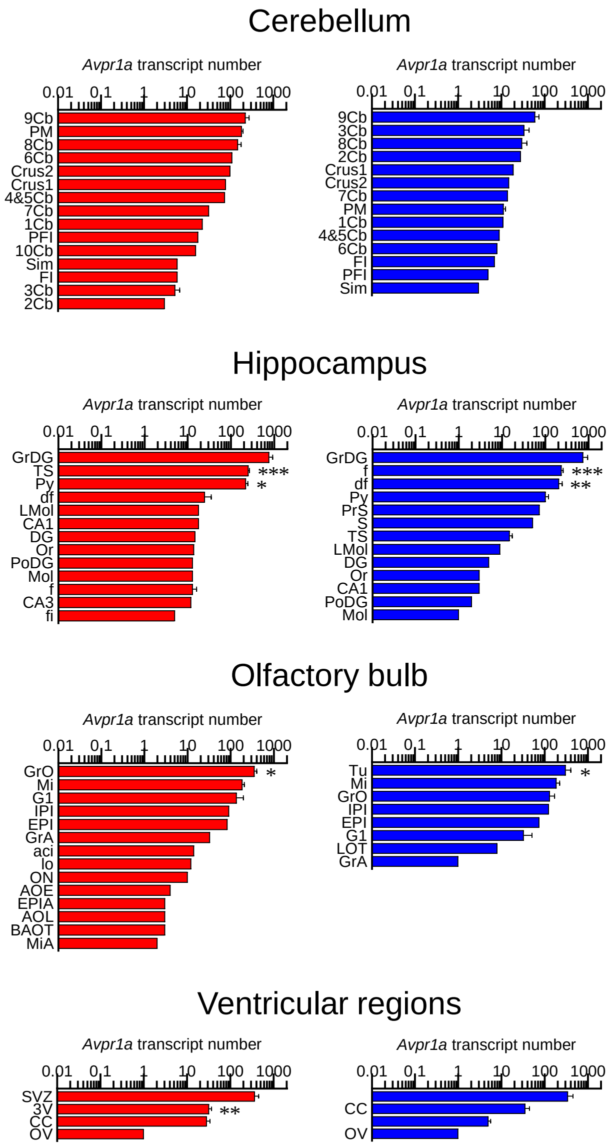
